# Development and validation of a next-gen health stratification engine to determine risk for multiple cardiovascular diseases

**DOI:** 10.1101/562900

**Authors:** Mehrdad Rezaee, Arsia Takeh, Igor Putrenko, Andrea Ganna, Erik Ingelsson

**Author notes:** Corresponding Author: (MR).

## Abstract

Cardiometabolic diseases (CMD) impose greater impact on every aspect of health care than any other disease group. Accurate and in-time risk assessment of individuals for their propensity to develop CMD events is one of the most critical paths in preventing these conditions. The principal objective of the present study is to report the development, and validation of a next generation risk engine to predict CMD. UK Biobank population data was used to derive predictive models for six CMD. Missing data were imputed using imputation algorithms. Cox proportional hazard models were used to estimate annual absolute risk and relative risk of different risk factors for these conditions. In addition to conventional risk factors, the applied model included socioeconomic data, lifestyle factors and comorbidities as predictors of outcomes. In total, 416,936 individuals were included in the analysis. The derived prediction models achieved consistent and moderate-to-high discrimination performance (C-index) for all diseases: coronary artery disease (0.79), hypertension (0.82), type 2 diabetes mellitus (0.87), stroke (0.79), deep vein thrombosis (0.75), and abdominal aortic aneurysm (0.90). These results were consistent across age groups (37-73 years) and showed similar predictive abilities amongst those with pre-existing diabetes or hypertension. Calibration of risk scores showed that there was moderate overestimation of CMD-related conditions only in the highest decile of risk scores for all models. In summary, the newly developed algorithms, based on Cox proportional models, resulted in high disclination and good calibration for several CMD. The integrations of these algorithms on a single platform may have direct clinical impact.

## Introduction

Cardiometabolic diseases (CMD) continue to be the leading causes of death in the United States since the 1920s, and 45% of the U.S. population is projected to suffer from any of these diseases by 2035 [1]. The healthcare cost associated with these diseases represent one of the greatest global economic burdens [2]. As with any chronic condition, appropriate prevention and selective treatment for CMD are the most effective approaches to defer their clinical and financial impact on individuals and across populations.

Primary prevention of chronic diseases is a resource intensive, costly, and non-effective if applied through non-selective implementation [3]. Therefore, accurate population and individual stratification is needed to provide individualized, as well as population-specific care. In order to achieve clinically relevant risk stratification, established risk factors and novel population-specific data should be considered to derive clinically applicable prediction algorithms.

For over 20 years, the concept of cardiovascular risk assessment has been tested through prediction models that are utilized in the clinical setting [4-6]. Current prediction models have good discrimination abilities to identify individuals who will develop CMD. However, there are opportunities to address the limitations of current models, such as inclusion of contemporary risk factors, biomarkers and genetic information as part of the algorithms [7]. Also, the currently systems are limited to only a few diseases, such as coronary artery disease and stroke, without consideration of major comorbidities. Moreover, current models do not allow for imputation for missing data; and finally, they are primarily directed to prevention of disease over a 10-year span. In this study, the development and validation of a next-gen stratification platform that integrates conventional clinical risk factors and biomarkers, socioeconomic, lifestyle factors and other co-morbidities data for six cardiometabolic diseases (CMD) is presented. To derive these new predictions models, we used data provided by the UK Biobank (UKBB) project [8], including over 400,000 men and women aged 37–73 years, with 6.1 years of median longitudinal follow-up.

## Materials and methods

### Baseline data preparation

Baseline data on 502,616 UKBB participants collected at assessment centers to derive the prediction models. Overall, 95% of the UKBB participants were self-described as white, with women comprising 54.4% of the total. CMD outcomes were determined based on International Classification of Diseases (ICD) edition 10 (ICD-10) codes, as well as self-reports for coronary artery disease (CAD), hypertension (HPT), type 2 diabetes mellitus (DM2), and deep vein thrombosis (DVT), and medications for CAD, HPT, and DM2. Six distinct datasets for each CMD were engineered. CAD was defined as I20–I25 and T82 codes. HPT was defined as I10, I15, and R03.0 codes. DM2 was defined as E11, E13, and E14 codes. Stroke was defined as G46.3, G46.4, I63, I66, I67, and I693 codes. DVT was defined as H34.8, H40.8, I23.6, I24.0, I63, I67.6, I74, I81, I82, I87.2, I87.3, K64.5, N48.8, N52.0, O03.3, O03.8, O04.8, O07.3, O08.7, 022, O87, Q26, T82.8, T83.8, T84.8, T85.8, and Z86.7 codes. Abdominal aortic aneurysm (AAA) was defined as I71 and I79.0 codes.

The UKBB data were subsequently linked to hospital episode statistics (HES) data from hospitals in England, Scotland and Wales. The age and date of a CMD event were determined based on primary or secondary ICD-10 codes in the HES data corresponding to the event using the earliest hospital record. The date of inclusion into the UKBB was defined as baseline and was used as starting point for time-to-event calculations. The exit date was determined as either date of death, end of follow-up (February 29, 2016), or a CMD event, whichever happened first. Only those CMD-positive cases that were identified by ICD-10 codes, self-reports, or medication as described above and had the date of the event determined based on the HES data were included into analyses, reducing the number of participants to 416,936. In addition, participants with prior CMD events (before baseline) were excluded from analyses of that specific event, e.g. those with prior CAD event were excluded from the CAD analyses and so on.

The datasets created for each CMD were spitted into training and testing sets based on 80%/20% ratio. Testing sets were used for model validation and calibration. Age- and CMD-specific testing sets were created by applying corresponding age and disease filters onto general test datasets (without reusing any data from the training sets to avoid overfitting).

### Variable definition

To develop highly predictive CMD risk prediction models, in addition to using already available UKBB data fields, the new variables were derived that captured sociodemographic and socioeconomic factors, laboratory test results, physiological measurements, physical activity, nutrition, alcohol consumption, family history of CMD; as well as the presence of diseases, disorders, or previous surgeries as shown in Table 1.

**Table 1.** Profile of variables for predicting the risk of six CMD.

|  | Type | N | 5% | 25% | 50% | 75% | 95% |
| --- | --- | --- | --- | --- | --- | --- | --- |
| Sex | binary | 416936 | women | women | women | men | men |
| BMI | continuous | 414265 | 20.84 | 23.94 | 26.48 | 29.58 | 35.82 |
| DBP | continuous | 415742 | 66 | 75 | 81 | 88 | 98.5 |
| Age | continuous | 416936 | 42 | 49 | 57 | 63 | 68 |
| FEV1 | continuous | 376770 | 60.83 | 82.19 | 93.79 | 104.38 | 119.88 |
| Current smoking | binary | 414793 | no | no | no | no | yes |
| Past smoking | binary | 412557 | no | no | yes | yes | yes |
| Family history of CAD | categorical | 359472 | no | no | no | yes (1)<sup>a</sup> | yes (2)<sup>b</sup> |
| Family history of DM2 | categorical | 385973 | no | no | no | no | mother |
| Family history of high blood pressure | categorical | 389301 | no | no | no | father | father and mother |
| Family history of stroke | categorical | 386630 | no | no | no | father | mother |
| Physical activity (MET x hours/week) | continuous | 379178 | 5.78 | 16 | 32 | 63 | 175.1 |
| Coffee consumption (cups) | continuous | 415021 | 0 | 0 | 2 | 3 | 6 |
| Alcohol score | continuous | 288169 | 0 | 0 | 2.5 | 10 | 10 |
| AHEI score | continuous | 199435 | 2.5 | 10 | 10 | 20.08 | 44.5 |
| Surgery history | binary | 322522 | no | no | no | no | yes |
| Hormone replacement therapy | categorical | 403518 | no | no | no | no | recent user (<3 years) |
| Hypercholesterolemia medication excluding aspirin | binary | 416936 | no | no | no | no | yes |
| Sleep apnea | binary | 416936 | no | no | no | no | no |
| Irritable bowel syndrome | binary | 416936 | no | no | no | no | no |
| Heart valve problem | binary | 416936 | no | no | no | no | no |
| Arrhythmia | binary | 416936 | no | no | no | no | no |
| Congestive heart failure | binary | 416936 | no | no | no | no | no |
| Hyperthyroidism | binary | 416936 | no | no | no | no | no |
| Education | categorical | 408500 | no | professional | professional | college or university | college or university |
| Income (£) | categorical | 353335 | <18,000 | 18,000 - 30,999 | 31,000 - 51,999 | 52,000 - 100,000 | >100,000 |
| Insomnia | categorical | 415605 | never/rarely | sometimes | sometimes | usually | usually |
| Sleep duration (hours) | categorical | 416117 | >4 and <6 or >9 and <11 | >=6 and <7 or >8 and <=9 | >=7 and <=8 | >=7 and <=9 | >=7 and <=10 |
| Lymphocyte | categorical | 395894 | >0.8 and <4.8 | >0.8 and <4.8 | >0.8 and <4.10 | >0.8 and <4.10 | >0.8 and <4.10 |
| Monocyte | categorical | 395894 | >0.2 and <0.9 | >0.2 and <0.9 | >0.2 and <0.9 | >0.2 and <0.9 | <=0.2 |
| MCH | categorical | 396632 | >=27 and <=34 | >=27 and <=34 | >=27 and <=34 | >=27 and <=34 | >34 |
| Platelet | categorical | 396631 | >=150 and <= 440 | >=150 and <= 440 | >=150 and <= 440 | >=150 and <= 440 | >=150 and <= 440 |
| RDW | categorical | 396633 | >= 11.6 and <= 14.6 | >= 11.6 and <= 14.6 | >= 11.6 and <= 14.6 | >= 11.6 and <= 14.6 | >14.6 |
| CAD age | continuous | 13607 | 45 | 54 | 58 | 62 | 67 |
| DM2 age | continuous | 8316 | 43 | 53 | 58 | 63 | 67 |
| HPT age | continuous | 36546 | 44 | 54 | 59 | 63 | 67 |
| DVT age | continuous | 7379 | 41 | 51 | 58 | 62 | 67 |

Types of variables, number of UKBB participants for each variable, and mean values (mode categories for categorical and binary variables) for different percentiles are shown. The number of participants for the CMD age variables corresponds to the number of prevalent cases.
<sup>a</sup>Either father, mother, or sibling
<sup>b</sup>Any combination of two of the following: father, mother, or sibling

Physical activity was assessed as the metabolic equivalent of task (MET) calculated in hours/week according to the “Guidelines for Data Processing and Analysis of the International Physical Activity Questionnaire (IPAQ) [9]. MET coefficients are indicated in Table 1. Alcohol score was calculated according to Alternative Healthy Eating Index (AHEI) guidelines [10]. One alcohol serving corresponded to 11.4 grams of alcohol. Further, a nutrition AHEI score was calculated as a sum of scores for the following nutrition categories: vegetables, fruits, grains, sugar sweetened beverages and fruit juices, nuts, meat, fish, PUFA, and alcohol. The nutrition scores were calculated according to AHEI guidelines [10].

In addition to the predicted CMD (target CMD), participants could of course experience other competing CMD outcomes. We used the age of experiencing these non-target diseases as an additional risk factor. For participants that did not experience a CMD event before baseline (CMD-negative cases), the age of CMD was set to 100. This approach allowed for incorporating time-dependent data without using the limitations of a modification of the Cox model, such as a Cox proportional hazards time varying model, which is often used to address time-dependency of predictors.

### Imputation of missing values

Multiple imputation by chained equations (MICE) implemented in Python (fancyimpute 0.3.1) and Bayesian ridge regression with the regularization parameter lambda of 0.001 was used for the imputation of missing values of continuous variables [11]. Parameters included initial filling with mean values, monotone visit sequence, the number of imputations = 100, the number of burn-in iterations = 10, no maximum and minimum possible imputed values, imputing with samples from posterior predictive distribution, the number of nearest neighbors for probabilistic moment matching = 5, and use of all columns to estimate current column. Cases with missing values in categorical variables were dropped before the imputation, and continuous variables were scaled to a range between 0 and 1.

### Variable selection for predictive modeling

Several approaches were employed for selecting variables included in the prediction model. Multicollinearity was first identified using pairwise correlation matrix (pandas 0.20.1), and the variables with the Pearson correlation coefficient higher than 0.3 were removed from the dataset. Recursive variable elimination with stratified 2-fold cross-validation (RFECV) on training datasets was then used to determine optimal number of variables by recursively considering smaller and smaller sets of variables (scikit-learn 0.20.0). One variable was removed at each iteration, minimum number of variables to be selected was one, and accuracy was used for scoring.

RFECV was used in combination with balanced random forest (imbalanced-learn 0.4.2) bivariate classification model. Parameters of the random forest model included the number of estimators = 100, Gini impurity as the quality of split, ‘auto’ sampling strategy, maximum depth of the decision tree = 0, minimum number of samples required to split an internal node = 2, minimum number of samples required to be at a leaf node = 1, minimum weighted fraction of the sum total of weights required to be at a leaf node = 0, the number of variables to consider when looking for the best split = ‘auto’, unlimited number of leaf nodes, minimum impurity decrease threshold for node splitting = 0, bootstrapping, random sampling without replacement, no use out-of-bag samples to estimate the generalization accuracy, the number of jobs to run in parallel for both fit and predict = 1, resampling all classes, but the minority class, the verbosity of the tree building process = 0, and balanced class weights.

In addition, principal component analysis (PCA) was used to validate the selection of variables and to avoid overfitting and poor calibration by determining that the number of selected variables is similar to the optimal number of principal components (scikit learn 0.20.0). The number of components to be retained was determined by using maximum-likelihood density estimation and full singular value decomposition (utilizing LAPACK library solver) as parameters of the PCA function, which applies Bayesian model selection to probabilistic PCA in this configuration [12].

### Predictive models and performance metrics

Linear Cox proportional hazard (PH) models and non-linear ensemble survival models were developed using lifelines 0.13.0 and scikit-survival 0.5 Python libraries, respectively. Two types of non-linear models were developed: decision tree-based gradient-boosting using Cox PH loss and gradient boosting with component-wise cubic smoothing splines as base learners.

Discriminative ability of the risk prediction models was assessed by Harrell’s concordance index (c-index) [13, 14, 15] calculated for testing datasets as the proportion of all comparable pairs in which the predictions and outcomes were concordant. Case pairs were comparable if at least one of them was CMD-positive. If the estimated risk was larger for the case with a lower time of event/censoring, the prediction of that pair was counted as concordant. If predictions were identical for a pair, 0.5 was added to the count of concordance. A pair was not comparable if an event occurred for both of them at the same time or an event occurred for one of them, but the time of censoring was smaller than the time of event of the first one. Prognostic indexes were used for the calculation of c-index.

In addition to c-index, we also used an additional metric for assessing the discriminative ability of Cox PH models, which was based on statistical ‘distance’ between the probabilities of experiencing a CMD event at certain time predicted for individuals from CMD-positive and CMD-negative groups. In the ‘distance’ approach, statistical significance of the difference between the two groups of probabilities was determined using one-way ANOVA. The result of this test was reported as an *F*-statistic with corresponding *p*-value.

Calibration of Cox PH models was evaluated by the Hosmer-Lemeshov goodness-of-fit test [16] and a calibration plot. The Hosmer-Lemeshow test was computed by partitioning the testing set into decile groups based on the predicted absolute risk of CMD events at time horizon of 5 years. Then, the number of CMD-positive and CMD-negative cases and the sum of the predicted probabilities for the both types of cases was calculated in each group as observed and not observed, and expected and not expected numbers, correspondingly. The Hosmer-Lemeshow test statistic was calculated using the following formula:

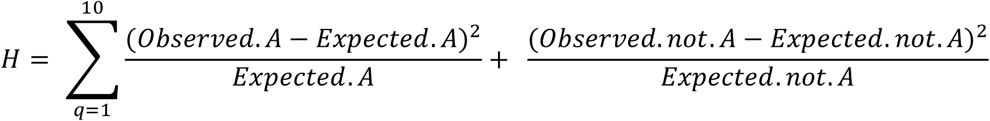

The resulted chi-square statistic was assessed using 8 degrees of freedom and was reported with *p*-value. A calibration plot was created by plotting the predicted risk probabilities against the observed risks for each group.

## Results

The study characteristics and the prevalence of six CMD at baseline for 416,936 UKB participants that include CMD-positive cases that were identified by ICD-10 codes, self-reports, or medication and had the date of the event determined based on the HES data are shown in Tables 1 and 2. Average age of men and women in this population was 56.3 ± 8.3 and 56 ± 8.1 years, correspondingly. During follow-up (median 6.1 years), 98,254 incident CMD events occurred in 67,785 participants that were free from the disease at baseline (Table 2).

**Table 2.** Prevalent and incident events for various CMD.

|  | Men |  | Women |  |
| --- | --- | --- | --- | --- |
|  | Prevalent events | Incident events | Prevalent events | Incident events |
| **CAD** | 9442 (5.11%) | 9560 (5.17%) | 4165 (1.79%) | 5479 (2.36%) |
| **HTN** | 19489 (10.54%) | 27939 (15.11%) | 17057 (7.35%) | 24724 (10.66%) |
| **DM2** | 5155 (2.79%) | 7590 (4.1%) | 3161 (1.36%) | 5209 (2.25%) |
| **Stroke** | 740 (0.4%) | 1866 (1.01%) | 446 (0.19%) | 1290 (0.56%) |
| **DVT** | 3870 (2.09%) | 7387 (4.0%) | 3509 (1.51%) | 6447 (2.78%) |
| **AAA** | 241 (0.13%) | 644 (0.35%) | 38 (0.016%) | 119 (0.051%) |

The prevalence of CMD at the baseline and incidence of CMD during the follow-up are shown in parenthesis.

### Imputation of missing data

Initial data quality evaluation showed that the number of missing values for examined variables (Table 1) varied from 0 to ∼52% with the mean of 6.3%, resulting in the no-null values dataset sizes of ∼78K – 81K (vs. initial ∼380K – 416K). As discussed in the methods, imputation of missing values for all continuous variables (Table 1) excluding CMD age variables, increased the sizes of CMD-specific datasets for predictive modeling to up to ∼195K – 215K. The discriminative ability of the CAD risk model trained on the imputed dataset with the sample size of 165,877 was tested on both imputed and unimputed datasets with the same sample size of 41,470 to validate the imputation. C-indexes calculated on the imputed and unimputed testing sets were 0.787 and 0.803, implying higher discriminative ability of the CAD model when tested on original, unimputed data.

### Predictive modeling

The discriminative ability of all Cox PH CMD models trained on the general population after the imputation of missing data varied between the diseases with highest and lowest c-indexes of 0.88 and 0.748 for AAA and DVT, respectively (Table 3). Cox PH models were further applied to calculate the risk probabilities of occurrence of a CMD event at 5 years following the initial observation. This time-to-event prediction was evaluated through determination of the statistical ‘distance’ between CMD-positive and CMD-negative test subgroups’ risk scores (Table 3). *F*-statistic values for the CMD models were highest for the models with high discriminative ability, except for the AAA model due to the low prevalence of this disease.

**Table 3:** Performance of CMD risk prediction models.

|  | C-index | Hosmer-Lemeshov test |  | ANOVA test |  |
| --- | --- | --- | --- | --- | --- |
|  |  | chi-2 | *p*-value | *F*-statistic | *p*-value |
| **CAD** | 0.787 | 55 | < 0.0001 | 24.7 | 1.80E-04 |
| **HPT** | 0.817 | 155 | < 0.0001 | 44.6 | 8.04E-07 |
| **DM2** | 0.873 | 54 | < 0.0001 | 36.6 | 1.20E-06 |
| **Stroke** | 0.783 | 18 | 0.02 | 17.6 | 6.20E-03 |
| **DVT** | 0.748 | 45 | < 0.0001 | 18.7 | 5.00E-03 |
| **AAA** | 0.88 | 17 | 0.03 | 15 | 1.20E-03 |

Performance is by c-index (discrimination), Hosmer-Lemeshov test (calibration), and the statistical ‘distance’ approach based on one-way ANOVA test (discrimination of risk probabilities). CMD-positive and negative groups were bootstrap sampled with replacement (N=100) to provide comparable *F*-statistic (*p*-values) across different disease endpoints.

Probability density function, which specifies the probability of predictions falling within a particular range of values for individuals from CMD-positive and CMD-negative test subgroups (Fig 1) was used for the visualization of the statistical ‘distance’ approach. The probability density function of the risk scores, as well as their distributions derived from different CMD models demonstrated that the range of risk scores for the CMD-positive subgroup was higher than that for the CMD-negative subgroup, and increased for CMD models characterized by higher c-index. Higher ratio between maximum values of the two probability density functions corresponded to higher discriminative ability.

**Fig 1.**
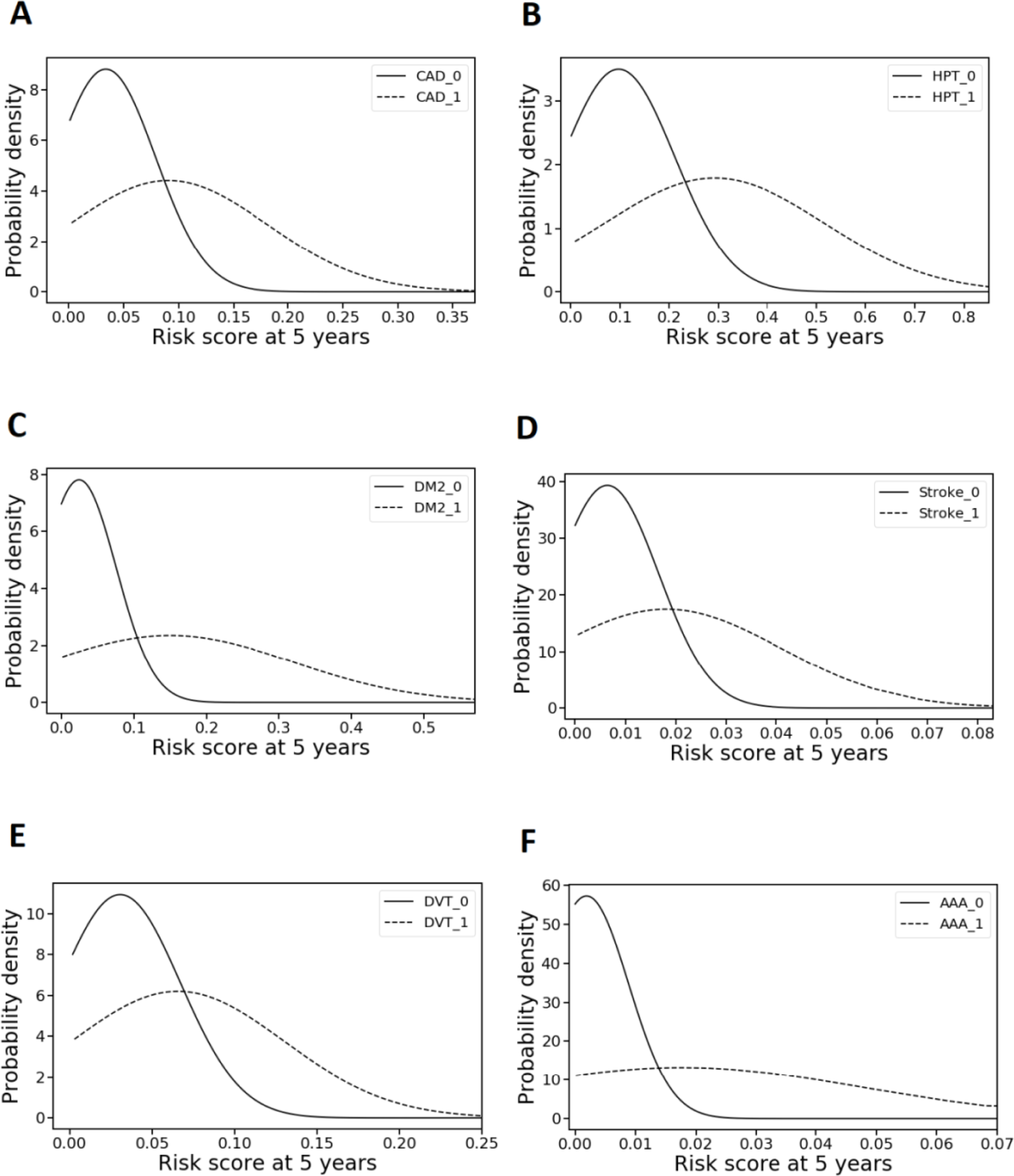
Statistical ‘distance’ approach. Probability density function expressed in relation to risk scores for six diseases (A-F) comparing participants developing CMD (CMD-positive,_1) and those who did not develop (CMD-negative, _0) within 5 years of follow up.

Assessment of the calibration properties for the CMD predictive models as calculated by the Hosmer-Lemeshow test (Table 3) and visualized by the calibration plot (Fig 2) showed adequate overall calibration, but moderate overestimation of CMD risk in the highest decile of risk scores.

**Fig 2.**
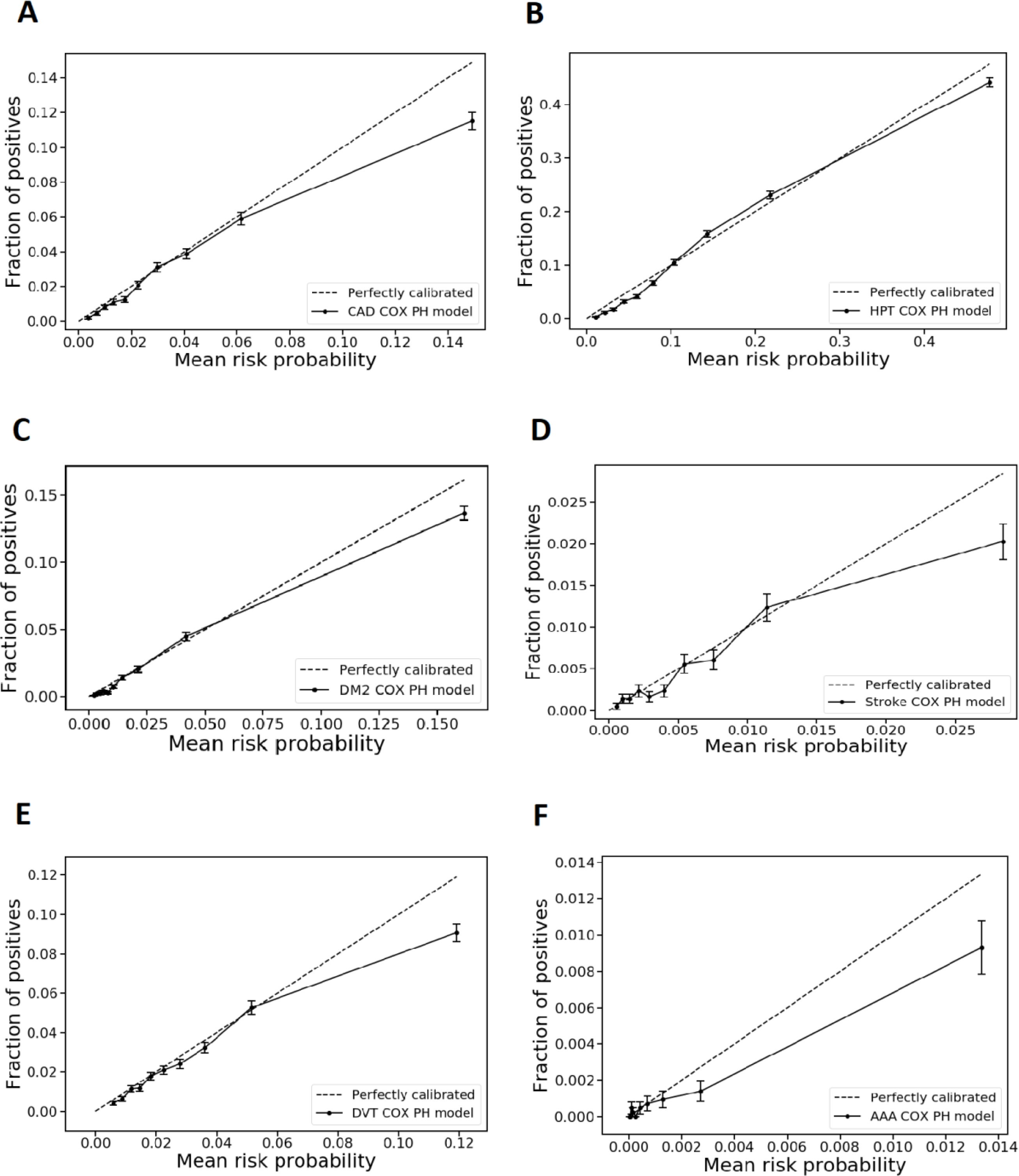
Calibration plots for CMD prediction models. Risk probabilities for six diseases (A-F), were split into deciles and mean risk probability for each decile was plotted vs. the portion of positive CMD cases in the decile for time horizon of 5 years.

In this study, the predictive performance of linear Cox PH models was compared with ensemble non-linear models as discussed in the methods. Non-linear survival models demonstrated comparable performance with the linear Cox model; however, this required significantly more computation time.

### CMD risk factors

To better understand the contribution of various risk factors to the pathophysiology of CMD, we ranked predictors of the risk of various CMD by the values of their regression coefficients (Table 4), indicating the degree of the association between the predictor and the outcome. Predictors presented in Table 4 represented only those with absolute values of coefficients larger than 0.8 and *p*-values less than 0.001 (see S1 Table for all coefficients). Statistical significance depended on the sample size and was affected by the prevalence of CMD. Accordingly, the number of predictors varied for each disease model.

**Table 4.**
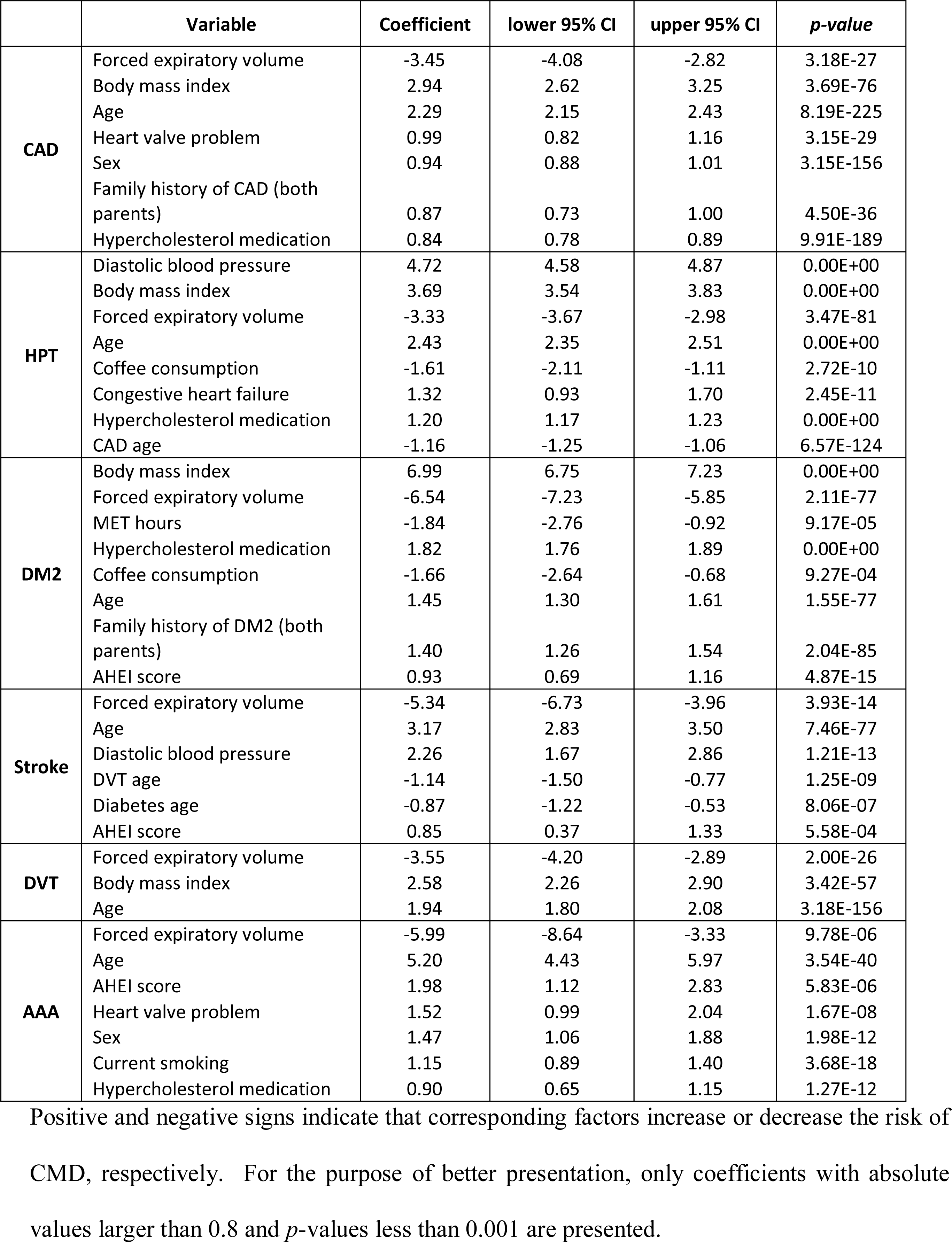
Ranked regression coefficients of predictors of the risk of various CMD models.

Across all disease models, age and low forced expiratory volume (FEV1) ranked as the most important predictors. Higher body mass index (BMI) and hypercholesterolemia medication were also among the strongest predictors for several models. Sex was ranked high only for the CAD and AAA, which is in a good agreement with our observation that the prevalence of these diseases was higher in men than in women. Family history ranked high only in predicting CAD and DM2. Nutrition was among the most important predictors for DM2, stroke, and AAA, which is likely explained by a healthier diet among individuals with certain risk factors and predispositions. Similarly, coffee consumption was an important predictor of HTN and DM2, possibly due to lower consumption in individuals with specific risk factor profiles. Physical activity was an important predictor only for DM2, and younger age of first occurrence of CAD, DVT and DM2 was among most important predictors for HTN and stroke, respectively.

### Validation

C-indexes for corresponding risk prediction benchmark models, with age and sex as the only predictors, were lower (delta, 0.04 – 0.2) when compared to those of our newly developed models. Broad range applicability and consistency of the performance of the developed risk prediction models for each disease were further determined by assessing the discriminative ability across subpopulations (Table 5). These subpopulations included (1) ‘healthy’ participants without any of the six target CMD at the baseline; (2) participants with at least one pre-existing non-target CMD at the baseline; and (3) various age categories. The performance of the models was highest in younger age and the healthy subgroup; while it significantly dropped in the subpopulation with pre-existing CMD.

**Table 5.**
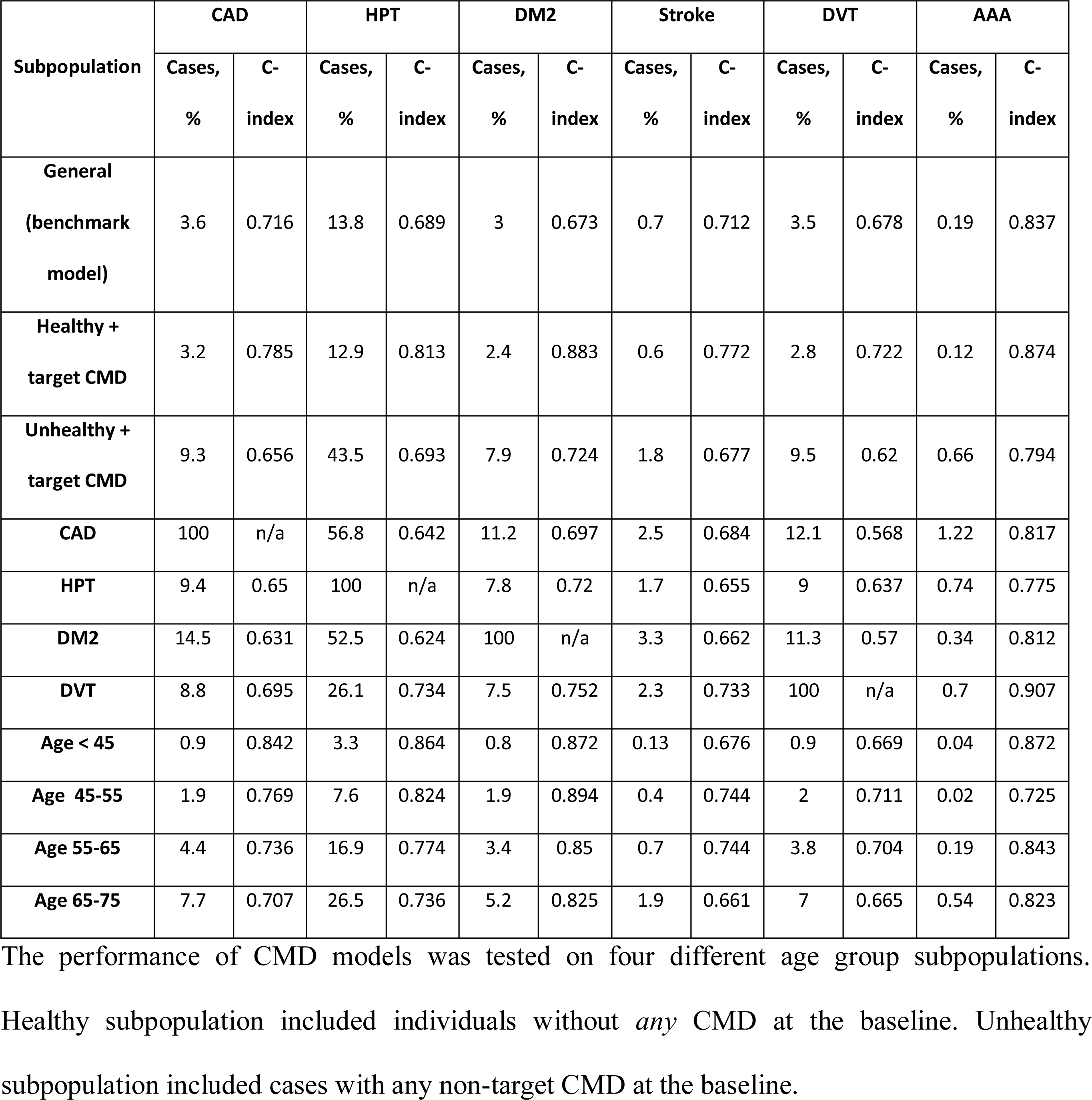
Validation of CMD risk prediction models.

## Discussion

### Principal findings

In this study, development and validation of a risk assessment platform applicable to six CMD is presented. The population-specific modeling for this platform was done using a dataset from the UK Biobank – a very large, longitudinal cohort study. This allowed us to derive prediction models and identify the most important contributing risk factors even for diseases with low incidence. Inclusion of a broad spectrum of risk factors allowed for modification of the array of input variables for the CMD risk prediction models included into the platform without significant decrease in their predictive performance. The models performed with high discriminative ability as demonstrated through extensive validation for different disease and age group subpopulations. Accordingly, this platform can accommodate different types of data sets and is applicable to population analysis, as well as individual assessment.

There is an abundance of risk predictors for CMD, and multiple prior attempts of combining them into risk calculators [17-19]. One of the major impediments for wide-spread application of these risk predictors includes lack of uniform validation through large population analyses. A comprehensive review found 363 models for cardiovascular risk stratification that have been developed and reported [20]. Only a minor collection of these models had sufficient evaluation according to contemporaneous analysis standards for either development or validation. For example, 39% of the 363 models analyzed utilized C-statistics for their development, and just over 60% for their validation. An even smaller number of the models utilized calibration as any part the performance measures. Although, the more recent models (since 2009) were more consistent in providing performance reports: 76% as part of their development, and up to 90% as part of validation [20].

In the current study, the discriminative ability of the developed models was similar or exceeded established models when available. For example, the Framingham Risk Score for coronary artery disease have been determined to be close to 0.76 and 0.79 for men and women, respectively [21]; these reported results were obtained only in the presence of all of the laboratory data and for a pre-selected small population. The modeling described for the platform in this report allows for incorporation of contemporary risk information. This is becoming increasingly important, since such more limited risk calculators may fail to express the accurate and true risk for a significant population. As demonstrated previously, either 50% of patients with CMD lack conventional risk factors or the conventional risk factors fail to explain more than 15-50% of the incidence of CHD [22-26].

The ability to incorporate socioeconomical data and nutritional information collectively can complement the basic information that is equivalent to conventional biomarkers. This is demonstrated in this study, as the performance of the current platform was achieved without the utilization of the blood laboratory information, such as lipid levels or blood glucose levels (as those were not available in UKBB at the time of this study). Utilization of a polygenic scoring is underway and can reveal a population at risk or protected from development of CMD [27-29]. It is expected that incorporation of the polygenic scoring will further increase the predicative performance of the current platform.

### Limitations of this study

Considering the fact that the UKBB population is not a complete representative of the UK or US populations, the main limitation of this study is that the developed models may need to be examined with inclusion of more diverse population. Predictive performance of the models was higher when tested on healthier and younger subpopulations. At the same time, training and calibration on CMD-specific datasets are required to improve discriminative ability of the models across CMD subpopulations. Considering the fact that the datasets used in predictive modeling were almost identical for different CMD, various predictive performances of the CMD models imply that despite overlapping pathophysiological pathways for various CMD, there are predictors specific for different CMD.

### Future directions

Considering computational limitations of non-linear survival models, bivariate time-dependent classification models utilizing machine learning algorithms can be used in future for determining the probability of CMD events at certain time horizons. The availability of relatively large healthcare datasets will further support the application of deep learning in time-dependent risk predictive modeling feasible. Incorporation of genetic and other-omics data may further improve the predictive functionality provided by this platform.

### Conclusions

In this report, we present development and validation of a new generation of disease risk prediction models. The differentiation variables of this platform include: a) assessment of multiple related diseases according to their associated outcomes (not just coronary artery disease); b) inclusion of contemporary risk factors; c) variable engineering and processing that allows for inclusion of data from different sources and addressing missing data points; d) population-specific stratification to assess risk prediction in different subgroups; e) being modular in nature to allow for inclusion of other risk determinants, such as genetic information; and f) being applicable at individual, as well as population level. These variables were designed into the platform in order to provide applicability of risk prediction to managing and changing the course of cardiometabolic diseases.

## Acknowledgements

This research has been conducted using the UK Biobank Resource under Application Number 24626.

## Supporting information

**S1 Table. Cox PH model regression coefficients for six CMD.** Regression coefficients (coef) and corresponding standard errors (se), p-values, lower and upper 95% confidence intervals are presented.

### Funding Source

The funder, Precision Wellness, Inc., provided support in the form of salaries for authors AT and IP, consultancy fees to AG, and as an unrestricted research grant to Stanford University (led by EI), but did not have any additional role in the study design, data collection and analysis, decision to publish, or preparation of the manuscript. MR did not receive any financial compensation for participation. The specific roles of these authors are articulated in the ‘Author contributions’ section.

### Author Contributions

1. Conceptualization: MR AT
2. Data curation: IP AT
3. Formal analysis: IP
4. Funding acquisition: MR
5. Investigation: MR AT EI
6. Methodology: MR AT IP AG EI
7. Project administration: MR AT
8. Resources: MR
9. Software: AT IP
10. Supervision: MR AT
11. Validation: MR EI AG
12. Visualization: IP
13. Writing – original draft: AT IP MR
14. Writing – review & editing: MR EI AG

